# An ancient fecundability-associated polymorphism switches a repressor into an enhancer of endometrial *TAP2* Expression

**DOI:** 10.1101/058388

**Authors:** Katelyn M. Mika, Vincent J. Lynch

## Abstract

Variation in female reproductive traits such as fertility, fecundity, and fecundability are heritable in humans, but identifying and functionally characterizing genetic variants associated with these traits has been challenging. Here we explore the functional significance and evolutionary history of a C/T polymorphism of SNP rs2071473, which we have previously shown is an eQTL for *TAP2* and significantly associated with fecundability (time to pregnancy). We replicated the association between rs2071473 genotype and *TAP2* expression using GTEx data and demonstrate that *TAP2* is expressed by decidual stromal cells at the maternal-fetal interface. Next, we show that rs2071473 is located within a progesterone responsive cis-regulatory element that functions as a repressor with the T allele and an enhancer with the C allele. Remarkably, we found this polymorphism arose before the divergence of modern and archaic humans, is segregating at intermediate to high frequencies across human populations, and has genetic signatures of long-term balancing selection. This variant has also previously been identified in GWA studies of immune related disease, suggesting both alleles are maintained due to antagonistic pleiotropy.

**Author Summary:** Female reproductive traits such as fertility and the time it takes to become pregnant are heritable. Many factors, including widespread contraceptive use and environmental influences, make identifying the genetic differences between individuals that are responsible for fertility differences between women difficult. We previously identified a common single nucleotide polymorphism that affects the expression of the gene *TAP2* and is significantly associated with how long it takes woman to become pregnant. Here we show that *TAP2* is expressed at the maternal-fetal interface in the uterus during pregnancy. We then show that the T version of the polymorphism functions to repress *TAP2* expression whereas the C form enhances *TAP2* expression. Remarkably, the C variant arose before the divergence of Neanderthals and modern humans and has become common in all human populations. This derived variant has previously associated with immune related diseases, suggesting the ancestral T and derived C variants are being maintained because they affect multiple traits.

## Introduction

Variation in female reproductive traits, such as age of menarche and menopause, fertility, age at first and last birth, fecundity, and fecundability are heritable in humans [1,2]. However, identifying the genetic bases for variation in most of these traits has proven challenging because of limited sample sizes, strong gene-environment interactions [1–3], widespread contraceptive use, and significant clinical heterogeneity among infertile couples. For example, while genome-wide association studies (GWAS) have identified loci associated with age of menarche and menopause [4–9], and age at first and last birth [10], few studies have successfully identified genetic variants associated with fertility, fecundity, and fecundability [3,10–12].

We recently performed an integrated expression quantitative trait locus (eQTL) mapping and association study to identify eQTLs in mid-secretory endometrium that influence pregnancy outcomes in a prospective study of Hutterite women [13–15]. Among the 189 eQTLs we identified, two were also associated with fecundability (time to pregnancy) [15]. The most significant association was rs2071473 (***P*** = 1.3×10^−4^), an eQTL associated with expression of ***TAP2*** in the HLA class II region. The C allele of rs2071473 was associated with longer intervals to pregnancy and higher expression of ***TAP2*** gene in mid-secretory phase (receptive) endometrium. The median time to pregnancy, for example, was 2.0, 3.1, and 4.0 months among women with the TT, CT, and CC genotypes, respectively [15].

An essential step in implantation is the establishment of receptivity by the hormone-primed endometrium. This ‘window of implantation’ occurs in the mid-secretory phase of the menstrual cycle [16], after endometrial stromal fibroblasts (ESFs) have differentiated (decidualized) into decidual stromal cells (DSCs) in response to progesterone and cyclic-AMP (cAMP) [17,18]. Decidualization underlies a suite of molecular, cellular, and physiological responses that support pregnancy including maternal immunotolerance of the fetal allograft [18,19]. Although function of TAP2 in the decidualized endometrium and the process of implantation have not been elucidated, TAP2 plays an integral role in translocating peptides from the cytosol to MHC class I molecules in the endoplasmic reticulum [20]. These data suggest that TAP2-dependent antigen processing and presentation by DSCs plays a role in establishing receptivity to implantation and immunotolerance at the maternal-fetal interface, likely by modifying the interactions between DSCs and immune cells in the decidualized endometrium [21].

Here we explore the functional significance and evolutionary history of the rs2071473 C/T polymorphism. We first replicate the association between rs2071473 genotype and ***TAP2*** expression using GTEx data and demonstrate that ***TAP2*** is expressed by DSCs at the maternal-fetal interface. Next, we show that rs2071473 is located within a cAMP/progesterone responsive regulatory element and disrupts a putative DDIT3 (CHOP10) biding site that functions as a repressor with the rs2071473 T allele and an enhancer with the C allele. Remarkably, we found that the C/T polymorphism arose before the divergence of modern and archaic humans, is segregating at intermediate to high frequencies across human populations, and has genetic signatures of long-term balancing selection.

## Methods

### rs2071473 is a multi-tissue eQTL for TAP2, HLA-DOB, and HLA-DRB6

We replicated the association between the T/C polymorphism at rs2071473 and ***TAP2*** expression levels using GTEx Analysis Release V6 (dbGaP Accession phs000424.v6.p1) data for 35 tissues including the uterus [22,23]. We also used GTEx data to identify other genes for which rs2071473 was an eQTL. Briefly, we queried the GTEx database using the ‘Single tissue eQTLs search form’ for SNP rs2071473 (http://www.gtexportal.org/home/eqtls/bySnp).

### *TAP2* is expressed by decidual stromal cells at the maternal-fetal interface

To determine the cell-type localization of TAP2 in the endometrium, we used data from the Human Protein Atlas immunohistochemistry collection (www.proteinatlas.org)[24] for endometrium, placenta, and decidua. We examined the expression of ***TAP2*** in the endometrium across the menstrual cycle [25], in mid-secretory phase endometrial biopsies from women not taking hormonal contraceptives (n=11), and women using either the progestin-based contraceptives depot medroxyprogesterone acetate (DMPA) or levonorgestrel intrauterine system (LNG-IUS) for at least 6 months [26], and in DSCs treated with a PGR-specific siRNA or a non-targeted siRNA [27] using previously generated microarray expression data. These microarray datasets were analyzed with the GEO2R analysis package (http://www.ncbi.nlm.nih.gov/geo/geo2r/), which implements the GEOquery [28] and limma R packages [29,30] from the Bioconductor project to quantify differential gene expression. We also examined ***TAP2*** expression in RNA-Seq data previously generated from ESFs treated with control media or differentiated (decidualized) with cAMP/MPA into DSCs [31,32].

### Cell culture and luciferase assays

#### Cell culture

Endometrial stromal fibroblasts (ATCC CRL-4003) immortalized with telomerase were maintained in phenol red free DMEM (Gibco) supplemented with 10% charcoal stripped fetal bovine serum (CSFBS; Gibco), 1× ITS (Gibco), 1% sodium pyruvate (Gibco), and 1% L-glutamine (Gibco). Decidualization was induced using DMEM with phenol red (Gibco), 2% CSFBS (Gibco), 1% sodium pyruvate (Gibco), 0.5mM 8-Br-cAMP (Sigma), and 1μM medroxyprogesterone acetate (Sigma).

#### Plasmid Transfection and Luciferase Assays

A 1000bp region spanning 50bp upstream of rs2071732 to the 3’-end of the PGR ChIP-Seq peak was cloned into the pGL3-Basic luciferase vector (Promega), once with the T allele and once with the C allele (Genscript). A pGL3-Basic plasmid without the 1kb rs2071473 insert was used as a negative expression control. DDIT3 and PGR expression plasmids were also obtained from Genscript. Confluent ESFs in 96 well plates in 80μl of Opti-MEM (Gibco) were transfected with 100ng of the luciferase plasmid, 100ng of DDIT3 and/or PGR as needed, and 10ng of pRL-null with 0.1 μl PLUS reagent (Invitrogen) and 0.25μl of Lipofectamine LTX (Invitrogen) in 20μl Opti-MEM. The cells incubated in the transfection mixture for 6hrs and the media was replaced with the phenol red free maintenance media overnight. Decidualization was then induced by incubating the cells in the decidualization media for 48hrs. After decidualization, Dual Luciferase Reporter Assays (Promega) were started by incubating the cells for 15mins in 20μl of 1× passive lysis buffer. Luciferase and renilla expression were then measured using the Glomax multi+ detection system (Promega). Luciferase expression values were standardized by the renilla expression values and background expression values as determined by pGL3-Basic expression.

### The rs2071473 C allele is derived in humans and segregating at intermediate to high frequencies across multiple human populations

To reconstruct the evolutionary history of the T/C polymorphism we used a region spanning 50bp upstream and downstream of rs2071473 from hg19 (chr6:32814778-32814878) as a query sequence to BLAT search the chimpanzee (CHIMP2.1.4), gorilla (gorGor3.1), organutan (PPYG2), gibbon, (Nleu1.0), rhesus monkey (MMUL_1), hamadryas baboon (Pham_1.0), olive baboon (Panu_2.0), vervet monkey (ChlSab1.0), marmoset (C_jacchus3.2.1), Bolivian squirrel monkey (SalBol1.0), tarsier (tarSyr1), mouse lemur (micMur1), and galago (OtoGar3) genomes. For all other non-human species we used the same 101 bp region as a query for SRA-BLAST against high-throughput sequencing reads deposited in SRA. The top scoring 100 reads were assembled into contigs using the ‘Map to reference’ option in Geneious v6.1.2 and the human sequence as a reference. Sequences for the Altai Neanderthal, Denisovan, Ust-Ishim, and two aboriginal Australians were obtained from the ‘Ancient Genome Browser’ (http://www.eva.mpg.de/neandertal/draft-neandertal-genome.html). The frequency of the C/T allele across the Human Genome Diversity Project (HGDP) populations was obtained from the ‘Geography of Genetic Variants Browser’ (http://popgen.uchicago.edu/ggv/).

We inferred ancestral sequences of the 101 bp region using the ancestral sequence reconstruction (ASR) module of the Datamonkey web-sever (http://www.datamonkey.org) [33] which implement joint, marginal, and sampled reconstruction methods [34], the nucleotide alignment of the 101 bp, the best fitting nucleotide substitution model (HKY85), a general discrete model of site-to-site rate variation with 3 rate classes, and the phylogeny shown in Fig 4A. All three ASR methods reconstructed the same sequence for the ancestral human sequence at 1.0 support.

### rs2071473 has signatures of balancing in humans and diversifying selection in primates

To infer if there was evidence for positive selection acting on the derived T allele we used an improved version of the EvoNC method [35] that has been modified into a branch-site model [36,37], and implemented in HyPhy v2.22 [37,38]. This method utilizes the MG94HKY85 nucleotide substitution model, which was also the best-fitting nucleotide substitution model for our target non-coding region and neutral rate proxy, and includes 10 replicate likelihood searches. Although this implementation of the EvoNC method that has been modified into a branch-site model capable of identifying positively selected sites in a priori defined lineages using Naive Empirical Bayes (NEB) and Bayes Empirical Bayes (BEB), previous studies have shown that these methods have high type-I and type-II error rates. Therefore, we inferred evidence for without reference to NEB or BEB identified sites and instead relied on a significant likelihood ratio test between the null and alternate models. We analyzed an alignment of the 101bp region described above, the phylogeny shown in Fig 4A, synonymous sites from the flanking ***HLA-DOB*** and ***TAP2*** genes which were identified from the same primate species using the method described above.

## Results

### rs2071473 is a multi-tissue eQTL for *TAP2*, *HLA-DOB*, and *HLA-DRB6*

We previously have shown that a T/C polymorphism at rs2071473 is an eQTL for ***TAP2*** in mid-secretory phase endometrium [15]. To replicate this observation in an independent cohort and in additional tissues, we tested if rs2071473 was correlated with ***TAP2*** expression using GTEx data. Similarly to our previous observation, we found that rs2071473 was an eQTL for ***TAP2*** in GTEx uterus samples (n=70, β=-1.1, ***P***=9.1×10^-11^; **Fig 1A**) as well as 34 other tissues (**Fig 1B**); the largest effect size was observed in the uterus (**Fig 1B**). We also used GTEx data to identify other genes for which rs2071473 was an eQTL and found that it was an eQTL for HLA-DOB in 25 tissues (β=−0.27 – −0.91, *P*=5.4×10^−9^ − 9.1 ×10^-20^; **Supplementary Table 1**) and for HLA-DRB6 in four tissues (β=0.37 − 0.46, *P*=6.7×10^-6^ – 2.1×10^-8^; **Supplementary Table 1**). However, rs2071473 was not identified as a uterine eQTL for HLA-DOB or HLA-DRB6 because neither gene is expressed in GTEx uterine tissues.

**Fig 1.**
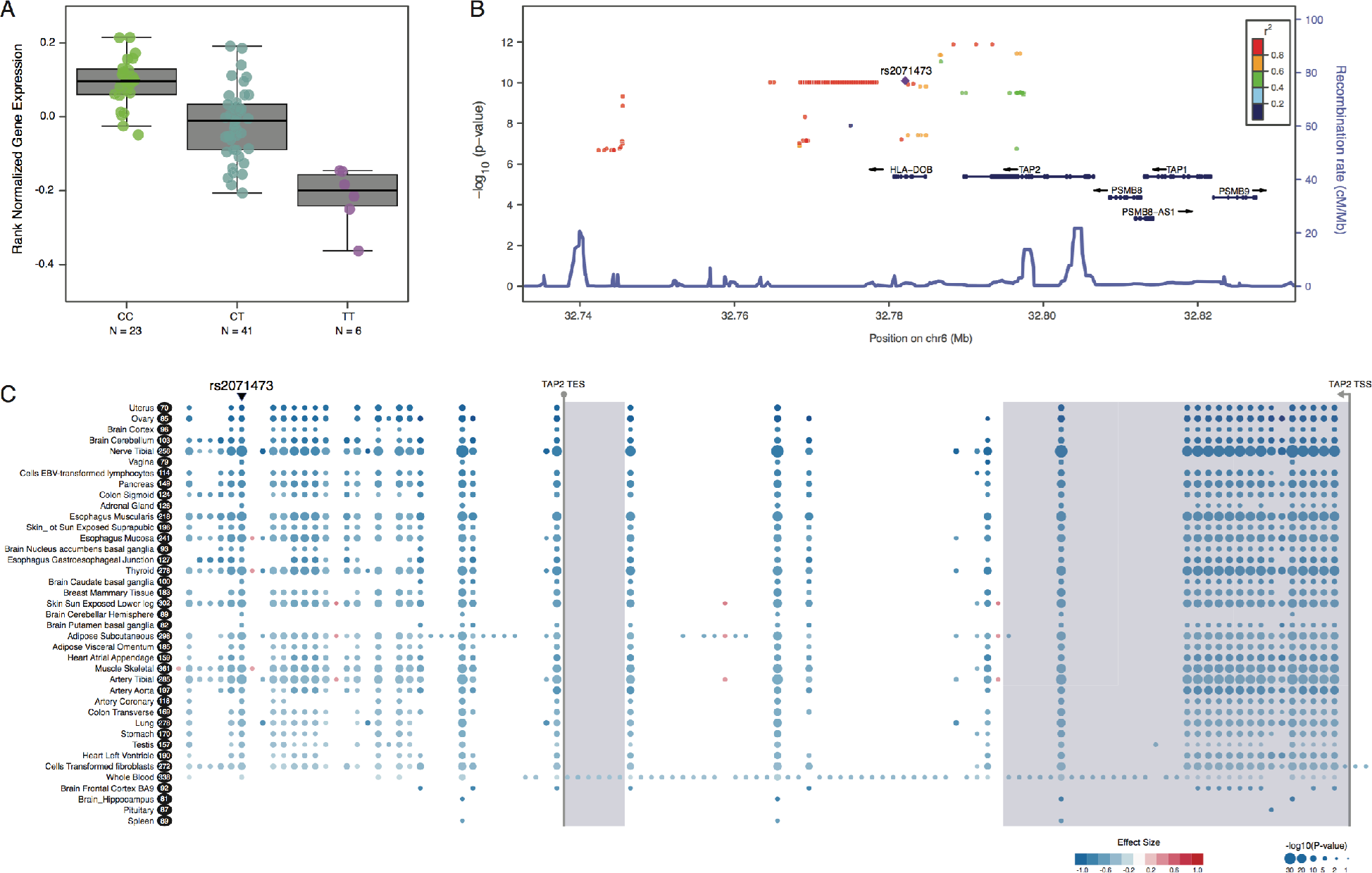
Replication of the rs2071473 C/T polymorphism as an eQTL for TAP2. (**A**) Uterine ***TAP2*** expression boxplots from GTEx data. Numbers below each boxplot are the number of individuals in each genotype. (**B**) Region association plot showing GTEx SNPs that are significant eQTLs for ***TAP2*** expression (−log_10_ P-value, left y-axis), color-coded base on their LD with rs2071473 (purple diamond). Local recombination rates are shown on the right y-axis). (**C**) Gene-eQTL Visualizer plot showing significant eQTLs for ***TAP2*** (see inset key for (−log_10_ P-value and effect size coding) across 39 GTEx tissues. Tissues are sorted by largest effect size top to bottom. Numbers in black ovals indicate sample sizes. The location of ***TAP2*** coding exons are shown with light blue shading, and the start site (TSS) and end site (TES) are indicated.

### *TAP2* is expressed by decidual stromal cells at the maternal-fetal interface

Although ***TAP2*** is expressed in the uterus, the mid-secretory phase endometrium is a complex tissue composed of numerous cell-types including perivascular mesenchymal stem-like cells [19,39], which are the likely precursors of endometrial stromal fibroblasts (ESFs)[18], decidual stromal cells (DSCs)[17,18], luminal and glandular epithelial cells, endothelial cells lining blood vessels, uterine natural killer cells (uNK)[40], uterine macrophage (uMP)[41,42], multiple populations of T-cells [43-46], and dendritic cells [47,48], among many others. To determine which cell-types express TAP2 we examined its localization in endometrial biopsies from the Human Protein Atlas immunohistochemistry collection. We found that TAP2 staining was localized primarily to the luminal and glandular epithelium and ESFs in non-pregnant endometrium (**Fig 2A**), was particularly intense in DSCs in pregnant endometrium (**Fig 2B**), and was absent from trophoblast cells (**Fig 2B/C**). Thus, TAP2 is expressed by maternal cells particularly DSCs at the maternal-fetal interface.

**Fig 2.**
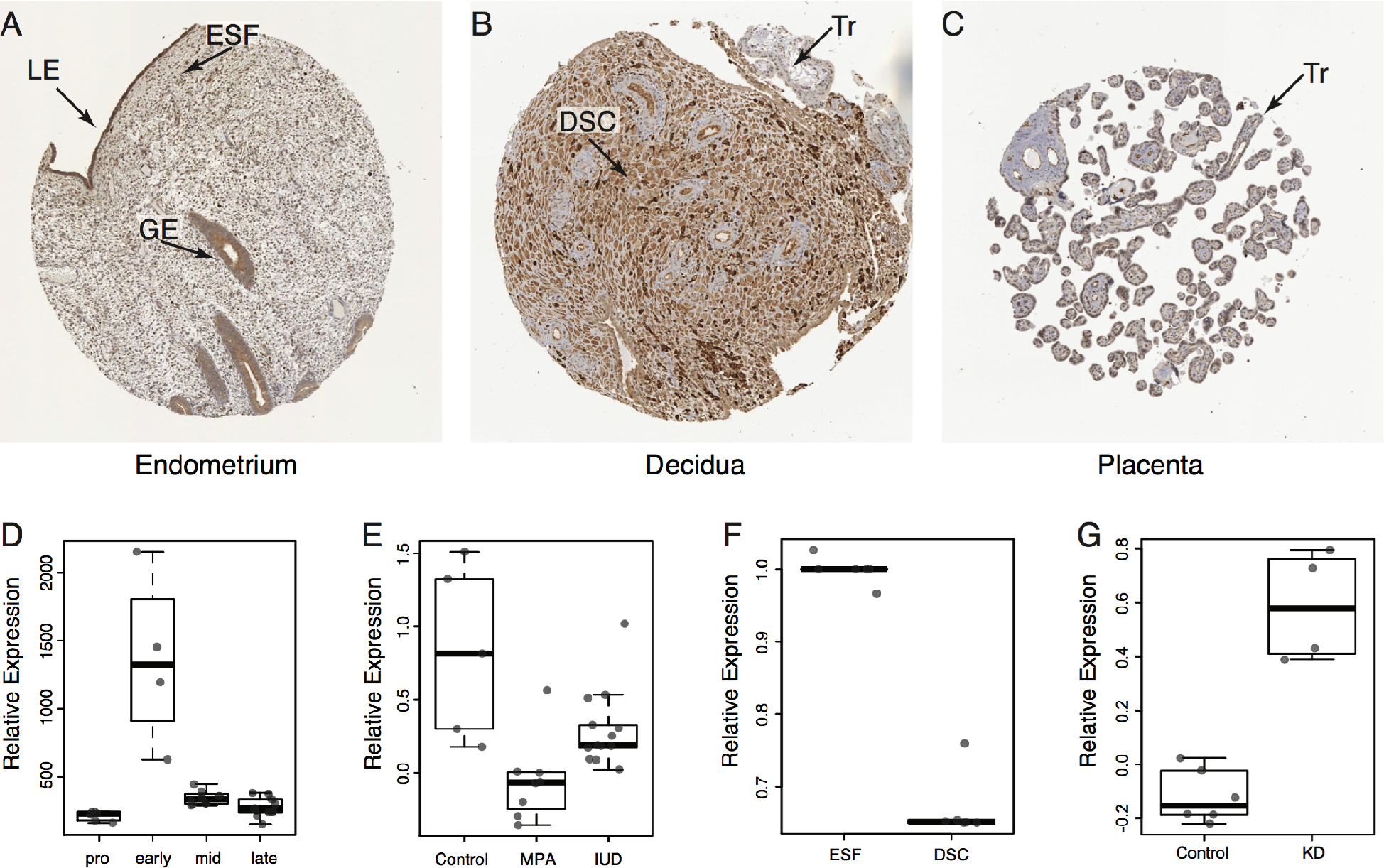
*TAP2* is regulated by progesterone and is expressed by decidual stromal cells at the maternal-fetal interface. (**A**) Immunohistochemistry staining showing TAP2 expression in the non-pregnant endometrium. LE, luminal epithelium. GE, glandular epithelium. ESF, endometrial stromal fibroblast. (**B**) Immunohistochemistry staining showing TAP2 expression in the pregnant endometrium (decidua). DSC, decidual stromal cell. Tr, trophoblast (placenta). (**C**) Immunohistochemistry staining showing TAP2 expression in the placenta. Tr, trophoblast. (**D**) Relative expression of ***TAP2*** across the menstrual cycle. (**E**) Relative expression of TAP2 in the endometrium of women using either the progestin-based contraceptives depot medroxyprogesterone acetate (MPA) or levonorgestrel intrauterine system (IUD). (**F**) Relative expression of ***TAP2*** in endometrial stromal fibroblasts treated with control media (ESFs) or differentiated into DSCs with cAMP/MPA for 48 hours. (**G**) Relative expression of ***TAP2*** in cAMP/MPA differentiated DSCs treated with a control siRNA or a PGR-specific siRNA (KD).

Next we tracked the expression of ***TAP2*** in the endometrium across the menstrual cycle using previously generated microarray expression data [25]. We found that ***TAP2*** expression reached its peak during the early secretory phase of the menstrual cycle and rapidly decreased in the mid-secretory phase (**Fig 2D**), suggesting that down-regulation of ***TAP2*** is regulated by progesterone and associated with endometrial receptivity to implantation. To infer if ***TAP2*** expression in the endometrium is regulated by progesterone we took advantage of an existing gene expression dataset of mid-secretory phase endometrial biopsies from women not taking hormonal contraceptives (n=11), and women using either the progestin-based contraceptives depot medroxyprogesterone acetate (DMPA) or levonorgestrel intrauterine system (LNG-IUS) for at least 6 months [26]. Consistent with regulation by progesterone, we found that ***TAP2*** expression was significantly lower in the endometria of women taking DMPA (***P***=0.002) or LNG-IUS (***P***=0.012) compared to controls (**Fig 2E**).

To directly test if ***TAP2*** is regulated by progesterone, we examined its expression in RNA-Seq data from ESFs treated with control media or differentiated (decidualized) with cAMP/MPA into DSCs [31,32] and found that ***TAP2*** was down-regulated ~33% by cAMP/MPA treatment (**Fig 2F**). To test if these effects were mediated by the progesterone receptor (PGR), we used a previously published dataset to compare ***TAP2*** expression in DSCs treated with a PGR-specific siRNA or a non-targeted siRNA [27] and found that knockdown of PGR significantly up-regulated TAP2 expression in DSCs (***P***=0.01). Thus we conclude that ***TAP2*** is down-regulated by progesterone during the differentiation of ESFs into DSCs and in the endometrium during the period of endometrial receptivity to implantation.

### The rs2071473 variant switches a repressor into an activator

Our observations that rs2071473 is an eQTL for ***TAP2*** and that ***TAP2*** expression is down-regulated by progesterone suggests rs2071473 may be located within or linked to a progesterone responsive enhancer. To identify such a regulatory element we used previously ChIP-Seq data from DSCs for the transcription factors PGR [32,49,50] and NR2F2 (COUP-TFII) [51], which regulate the transcriptional response to progesterone and immune genes, respectively, H3K27ac which marks active enhancers, H3K4me3 which marks active promoters [32], and DNaseI-Seq and FAIRE-Seq to identify regions of open chromatin [32]. We found that rs2071473 was located 260bp upstream of PGR and NR2F2 binding sites, within local DNase FAIRE peaks, and in a region of elevated H3K4me3 and H3K27ac signal (**Fig 3A**) suggesting this region is a progesterone responsive c/s-regulatory element.

**Fig 3.**
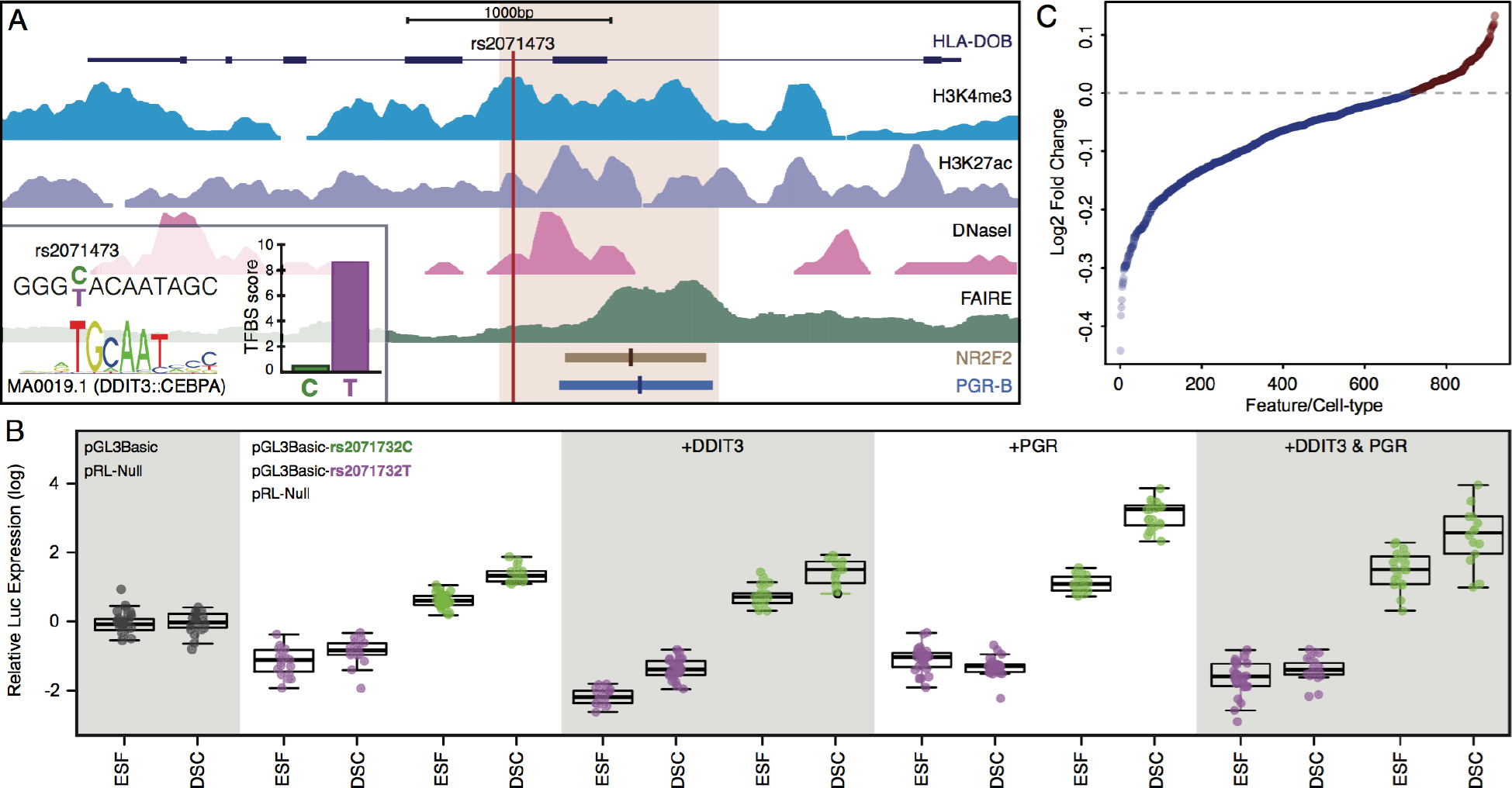
The C/T polymorphism at rs2071473 is located within a progesterone responsive c/s-regulatory element in decidual stromal cells. (**A**) Location of rs2071473 with respect to histone modifications that characterize promoters (H3K4me4 ChIP-Seq), enhancers (H3K27ac ChIP-Seq), open chromatin (DNaseI-Seq, FAIRE-Seq), as well as PGR and NR2F2 ChIP-Seq binding sites. Inset, the rs2071473-C allele is predicted to disrupt a DDIT3 binding site. The 1kb region shown in light brown was cloned into the pGL3Basic luciferase reporter vector for functional characterization. (**B**) Luciferase assay results testing the regulatory potential of the T (pGL3Basic-rs2071473C) and C (pGL3Basic-rs2071473T) alleles in endometrial stromal fibroblasts treated with control media (ESFs) or differentiated into DSCs with cAMP/MPA for 48 hours. Data are shown as luciferase expression from the pGL3Basic-rs2071473C or pGL3Basic-rs2071473T reporter relative to renilla expression (pRL-null and empty vector (pGL3Basic) controls. +DDIT3, Relative luciferase expression from the pGL3Basic-rs2071473C or pGL3Basic-rs2071473T in ESFs and DSCs co-transfected with DDIT3. +PGR, Relative luciferase expression from the pGL3Basic-rs2071473C or pGL3Basic-rs2071473T in ESFs and DSCs co-transfected with PGR. +DDIT3 & PGR, Relative luciferase expression from the pGL3Basic-rs2071473C or pGL3Basic-rs2071473T in ESFs and DSCs co-transfected with DDIT3 and PGR. (**C**) DeepSea predicted effects (log_2_ fold change) of rs2071473 on regulatory features across cell-types.

To test if this locus has regulatory potential we synthesized a 1000bp region spanning 50bp upstream of rs2071732 to the 3’-end of the PGR ChIP-Seq peak (**Fig 3A**) with either the reference C allele or the alternate T allele and cloned them into the pGL3-Basic luciferase reporter vector, which lacks both an endogenous promoter and enhancer. Next we transiently transfected either the pGL3Basic-rs2071732T or pGL3Basic-rs2071732C luciferase reporter along with the pRL-null internal control vector into ESFs and DSCs and quantified luciferase and renilla expression using a dual luciferase assay.

We found that luciferase expression was significantly lower in ESFs (Wilcox test ***P***=1.75×10^−11^) and DSCs (Wilcox test ***P***=2.87×10^−9^) transfected with the pGL3Basic-rs2071732T reporter compared to empty pGL3Basic vector controls (**Fig 3B**). In stark contrast, luciferase expression was significantly higher in ESFs (Wilcox test ***P***=1.42×10^−9^) and DSCs (Wilcox test ***P***=2.53×10^−13^) transfected with the pGL3Basic-rs2071732T reporter compared to controls (**Fig 3B**). The difference in luciferase expression between the pGL3Basic-rs2071732T and pGL3Basic-rs2071732C was also significant in ESFs (Wilcox test ***P***=2.50×10^−12^) and DSCs (Wilcox test ***P***=1.76×10^−11^). Luciferase expression was significantly induced upon differentiation of ESFs to DSCs by cAMP/MPA treatment in both the pGL3Basic-rs2071732T (Wilcox test ***P***=0.02) and pGL3Basic-rs2071732C (Wilcox test ***P***=2.53×10^−13^) transfected cells (**Fig 3B**). Thus we conclude that the locus in which rs2071473 resides is a progesterone responsive *cis*-regulatory element and that the T/C polymorphism switches an enhancer into a repressor.

### The rs2071473 C allele likely disrupts a DDIT3 binding site

The T/C polymorphism at rs2071473 is several hundred base pairs away from the PGR and NR2F2 binding sites and is therefore unlikely to directly effect their binding (**Fig 3A**). However our luciferase assay results indicate that the T/C polymorphism has regulatory effects, suggesting this polymorphism disrupts a binding site for a transcriptional repressor or creates a binding site transcriptional activator. To infer which of these scenarios was most likely we identified putative transcription factor binding sites in a 25bp window upstream and downstream of rs2071473 using ConSite [52] and JASPAR transcription factor binding site profiles [53]. We found that the T/C polymorphism occurs at an invariant T site in the DDIT3 motif (TGCAAT), which was predicted to abolish DDIT3 binding (**Fig 3A**, inset). Similarly, the T/C polymorphism was predicted by DeepSea [54], a deep learning-based algorithm that infers the effects of single nucleotide substitutions on chromatin features such as transcription factors binding, DNase I sensitivitity, and histone marks, to generally have a negative effects on regulatory functions across multiple cell-types (**Fig 3C**).

While DDIT3 (also known as CH0P10 and GADD153) was initially characterized as a dominant negative inhibitor of CEBP family transcription factors [55], it can also function as a transcription factor [56,57] and is transcriptionally regulated the cAMP [58]. Indeed we found that CH0P10 expression was down-regulated by cAMP/MPA in our RNA-Seq data from ESFs (TPM=187.68) and DSCs (TPM=61.20). These data suggest that the T/C polymorphism may disrupt a DDIT3 binding site that mediates transcriptional repression of ***TAP2***, unmasking a secondary enhancer function. To test this hypothesis we co-transfected ESFs and DSCs with a DDIT3 expression vector, the pRL-null internal control vector, and either the pGL3Basic-rs2071732T or pGL3Basic-rs2071732C luciferase reporter, and compared luciferase expression to control ESFs and DSCs. If DDIT3 mediates repression by biding the T allele, then co-transfection of DDIT3 with the pGL3Basic-rs2071732T reporter should augment repression whereas co-transfection with the pGL3Basic-rs2071732C reporter should be unaffected. Indeed, expression of DDIT3 significantly reduced luciferase expression from the pGL3Basic-rs2071732T reporter in ESFs (Wilcox test ***P***=3.48×10^-10^) and DSCs (Wilcox test ***P***=1.591e-05) but had no effect on luciferase expression from the the pGL3Basic-rs2071732C reporter in either ESFs or DSCs (P> 0.21; **Fig 3B**).

To test if this regulatory element is progesterone-responsive, we co-transfected ESFs and DSCs with a PGR rather than a DDIT3 expression vector and repeated the luciferase assays described above. Expression of PGR did not affect luciferase expression from the pGL3Basic-rs2071732T reporter in ESFs (Wilcox test ***P***=0.67) but did enhance repression in DSCs (Wilcox test ***P***=9.56×10^−5^; **Fig 3B**). Co-transfection of PGR elevated luciferase expression from the pGL3Basic-rs2071732C reporter in both ESFs (Wilcox test ***P***=2.82×10^−8^) and DSCs (Wilcox test ***P***=2.53×10^−13^; **Fig 3B**). Finally we tested whether DDIT3 and PGR acted cooperatively or antagonistically by co-transfecting both the DDIT3 and PGR expression vectors with either the pGL3Basic-rs2071732T or pGL3Basic-rs2071732C reporters into ESFs and DSCs. Consistent with a weakly antagonistic functional interaction, luciferase expression in DDIT3 and PGR transfected ESFs and DSCs was intermediate between luciferase expression DDIT3 or PGR transfected cells (**Fig 3B**). These data suggest that the reference C allele likely disrupts a DDIT3 biding site, which unmasks a progesterone responsive enhancer.

### The rs2071473 C allele is derived in humans and has evidence of positive selection

To reconstruct the evolutionary history of the T/C polymorphism we identified a region spanning 50bp upstream and downstream of rs2071473 from 45 primates, including species from each the major primate lineages, multiple sub-species of African apes (Homininae), as well as modern and archaic (Altai Neanderthal and Denisova) humans. Next we used maximum likelihood methods to reconstruct ancestral sequences for this 101 bp region. We found that the T allele was ancestral in primates and that the C variant was only found in modern human populations as well as the Altai Neaderthal genome (**Fig 4A**). Next we examined the frequency of these alleles across the Human Genome Diversity Project (HGDP) populations as well as two aboriginal Australians and found that the derived and ancestral alleles were segregating at intermediate to high frequencies in nearly all human populations (**Fig 4B**). These results indicate that the derived C variant at rs2071473 arose before the divergence of modern and archaic human lineages.

**Fig 4.**
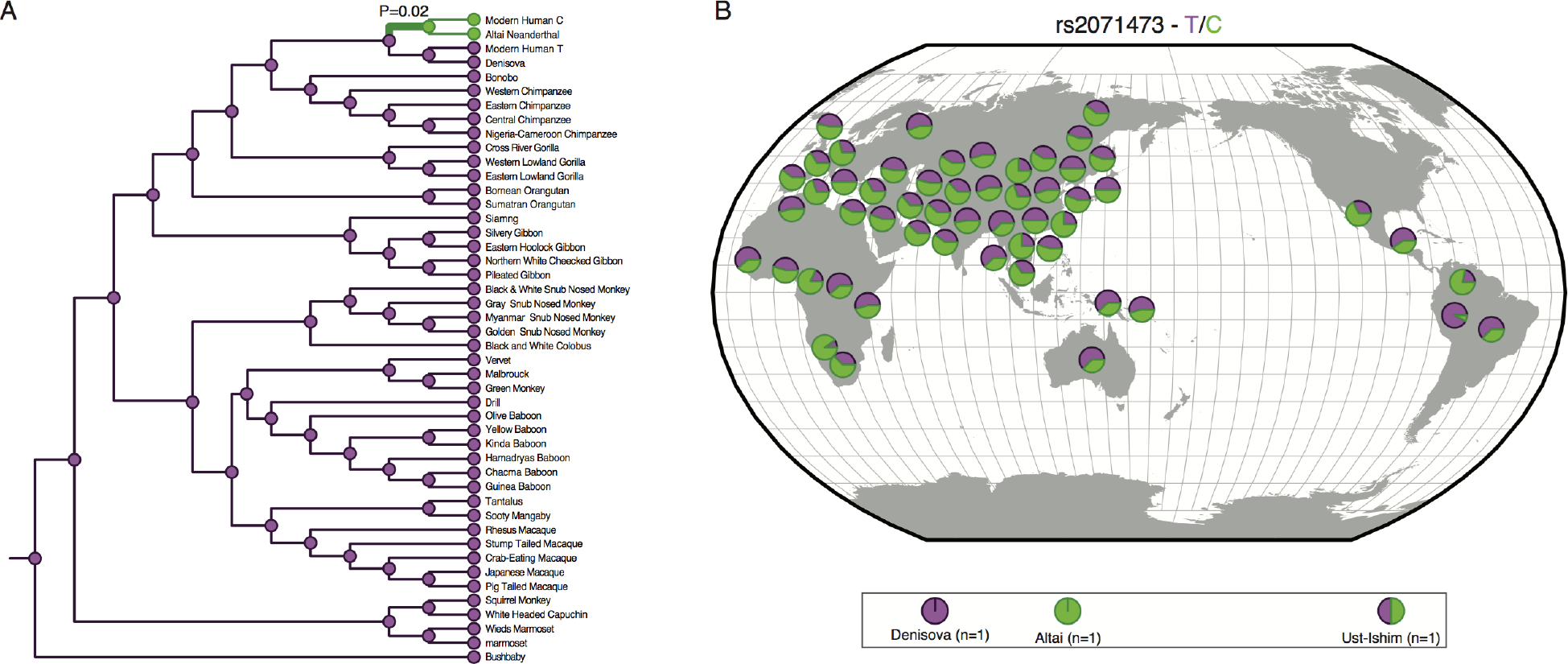
Evolutionary history C/T polymorphism at rs2071473. (**A**) rs2071473 genotype across extant and ancestral primates. Genotype in extant species is shown as circles next to species names at terminal branches. Genotype at internal nodes based on ancestral reconstruction is shown as circles. Purple, T. Green, C. ***P***=0.02, indicates lineage identified as positive selected based on the branch-sites version of the EvoNC method. (**B**) Distribution of the T and C alleles of rs2071473 across HGDP populations.

To test if the derived C allele may have been positively selected we used a branch-sites version of the EvoNC method [35–37]. In this analysis, the nucleotide substitution rate in noncoding regions (*d_nc_*) is compared with a neutral rate proxy from either introns or synonymous substitutions (*d_S_*) in nearby coding genes. The strength and direction of selection acting on noncoding regions is given by *d_nc_*/*d_S_* or ζ, which is analogous to the *d_N_*/*d_S_* rate or ω, with ζ = 1 indicating neutral evolution, ζ < 1 indicating negative (purifying) selection, and ζ > 1 indicating positive selection. We analyzed the 101 bp region described above and synonymous sites from the flanking HLA-DOB and TAP2 genes, and fit three models to the data: 1) A null model that constrains ζ≤ 1 across all sites and lineages; 2) An alternate model that allows for ζ ≤ 1 in foreground and background branches, ζ ≤ 1 in background branches and ζ > 1 in the foreground branch; and 3) An alternate model that allows for ζ < 1 in foreground and background branches, ζ = 1 in foreground and background branches, ζ < 1 in background and ζ > 1 in foreground branches, and ζ = 1 in background and ζ > 1 in foreground branches. The null model was rejected in favor of both alternate model 1 (alternate model 1 LRT=5.37; ***P***=0.02) and alternate model 2 (alternate model 2 LRT=5.38; ***P***=0.02), suggesting the T to C substitution may have been positively selected.

### rs2071473 has signatures of balancing in humans and diversifying selection in primates

Our observation that the C allele originated before the divergence of modern and archaic humans and is segregating at intermediate frequencies across modern human populations suggests that the ancestral and derived variants may be maintained by balancing selection. Previous studies have shown that balancing selection is common in the HLA region in which ***TAP2*** and rs2071473 are located [59–62]. DeGiorgio et al, for example, developed a model-based approach to identify signatures of ancient balancing selection and found that the *HLA-DOB* lcous, in which rs2071473 is located, was an outlier (top 0.5% of all scores across the genome) in their scan for balancing selection [63]. Consistent with the action of long-term balancing selection, rs2071473 has essentially no differentiation between populations (F_st_ = 0 – 0. 016), and is located in region with a relative excess of common polymorphisms across CEU and YRI populations as measured by Tajima’s D (1.16-2.07), Fay and Wu’s H (−23.16 – −56.83), and Pi (86.41-107.71)(**Fig 5**).

**Fig 5.**
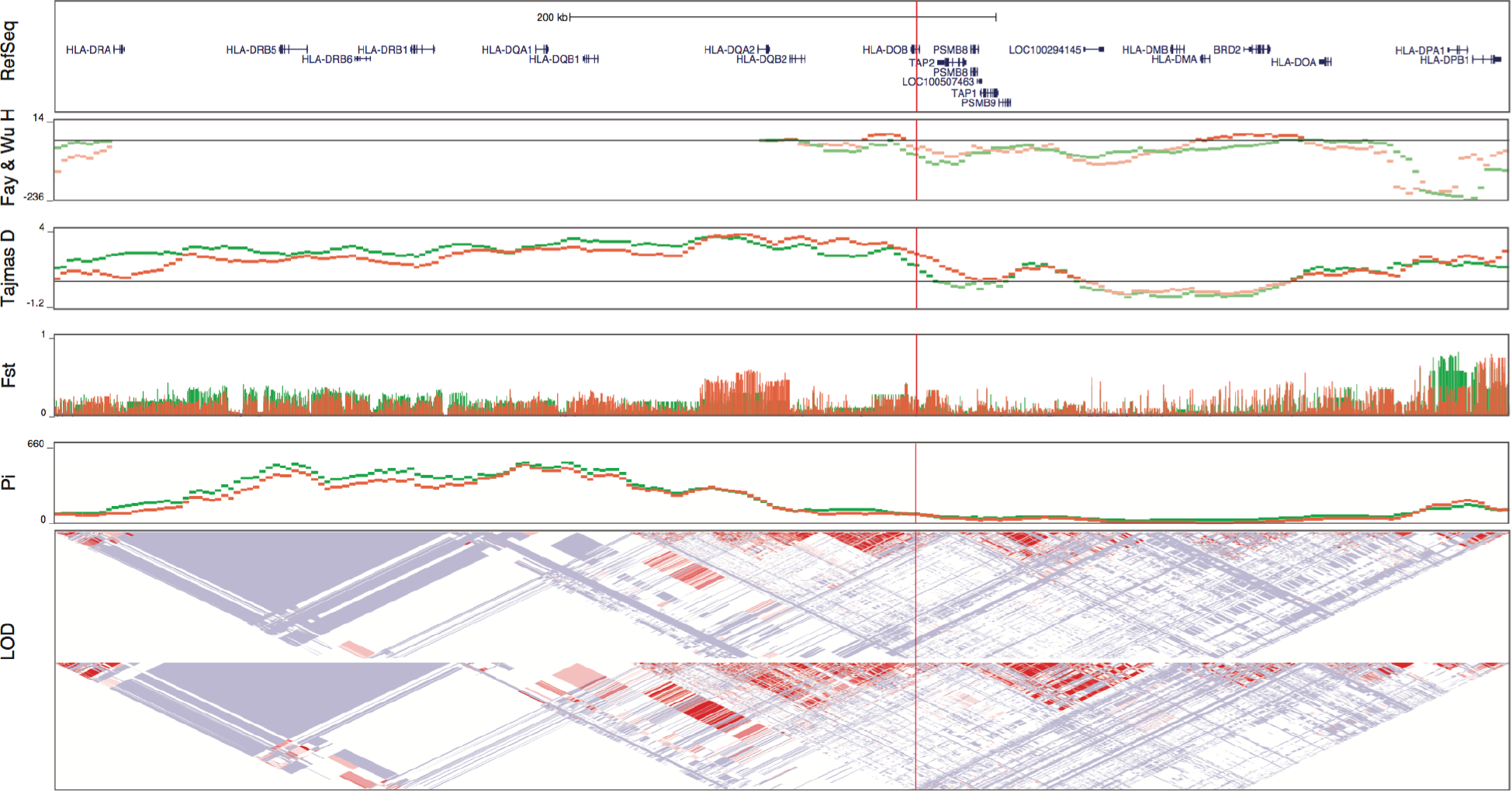
The rs2071473 C/T polymorphism has genetic signatures of balancing selection. Fay and Wu’s H statistic, Tajima’s D, Fst, and nucleotide diversity (Pi) for HGDP CEU (green) and YRI (red) populations across the HLA-region of chromosome 6. The location of rs2071476 is shown with a vertical red line. Linkage disequilibrium across this region is shown the log odds score (LOD) with white diamonds indicate pairwise D’ values less than 1 with no statistically significant evidence of LD (LOD < 2), light blue diamonds indicate high D’ values (>0.99) with low statistical significance (LOD < 2), and light pink diamonds indicate high statistical significance (LOD >= 2) but the low D’ (less than 0.5).

To test if in the rs2071473 enhancer region evolve under positive diversifying selection across primates we used the sites version of the EvoNC method [35–37]. We fit three models to the alignment described above: 1) A null model that constrains ζ ≤ 1 across all sites; 2) An alternate model that allows for categories of sites with ζ ≥ 1 and ζ < 1; and 3) An alternate model that allows for categories of sites with ζ < 1, ζ = 1, or ζ > 1. We found that the null model was not rejected in favor of alternate model 1 (LRT=0; ***P***=1), however, alternative model 2 was a better fit to the data than either the null model LRT=11.76; ***P***=0.019) or alternative model 1 (LRT=11.78; ***P***=0.0005). These data indicate that including distinct rate classes for sites with ζ < 1 or ζ = 1 significantly improves the alternate model, and suggest that positive diversifying selection acts on sites in this regulatory element across primates.

## Discussion

The mechanisms that promote maternal tolerance of the antigentically distinct fetus are complex [64] and have been the subject of intense study since Medawar formulated the immunological paradox posed by pregnancy and proposed tolerance was achieved by physical separation of maternal and fetal tissues, maternal immunosuppression, and immaturity of fetal antigens [65]. It is now clear that rather than being a site of maternal immunosuppression [66], the maternal immune system in the endometrium plays an active role in establishing a permissive environment for implantation, placentation, and gestation. For example, maternal regulatory T cells (Tregs) [44,67], shifts in the Th1/Th2/Th17 balance [45], uterine natural killer cells [40,68], uterine dendritic cells [47], uterine macrophage [69], and signaling by DSCs [70–72] all contribute to establishing immunotolerance and even promote placental invasion into maternal tissues.

It has also become clear that the major histocompatibility complex (MHC) genes, which play an important role in the rejection of non-self tissues, contribute to maternal tolerance of the fetus [21]. Matching of HLA antigens between couples, for example, is associated with longer intervals from marriage to birth compared to couples not matching for HLA [73,74]; these longer intervals result from both higher miscarriage rates among couples matching for class I HLA-B antigens [14] and longer intervals to pregnancy among couples matching for class II HLA-DR antigens [13]. Similarly, the non-classical HLA class I genes HLA-F are associated with fecundability whereas overwhelming evidence indicates maternal and fetal HLA-G genotypes are associated with miscarriage, recurrent pregnancy loss, and preeclampsia [75–86]. Collectively these data implicate antigen presentation by MHC as one of the molecular mechanisms that underlie successful implantation, maternal immunotolerance, and the establishment and maintenance of pregnancy.

The ATP-binding cassette transporter TAP, a heterodimer composed of TAP1 and TAP2, translocates peptides from the cytosol to awaiting MHC class I molecules in the endoplasmic reticulum, which results in cell surface presentation of the trimeric MHC complex to immune cells such as T lymphocytes and natural killer cells [20]. Loss and reduced TAP expression leads to loss and reduced surface HLA expression [87–89], altered surface HLA repertoires [90], and the surface expression of distinct antigenic peptides that are recognized by cytotoxic T lymphocytes [91]. Our observation that TAP2 is highly expressed by DSCs at the maternal-fetal interface and that ***TAP2*** expression levels are associated with fecundability suggests that changes in TAP2 stoichiometry may alter MHC processing and thus interactions between DSCs and maternal immune cells. Indeed DSCs express numerous HLA class I molecules, including HLA-G and HLA-F [92,93], which are transcriptionally up-regulated by progesterone during decidualization [72].

While the connection between TAP2 expression levels in DSCs and immune signaling are obvious, our observation that the ancestral and derived alleles of rs2071473 arose before the divergence of modern and archaic humans and has signatures of long-term balancing selection is unexpected. The HLA region has long been recognized to be under balancing selection [59,60,63,94,95]. These signals, however, are usually attributed to polymorphisms within the classical HLA class I genes rather regulatory regions (cf. [96]). TAP2 and rs2071473 are also located near the distal end of the HLA region and bounded by regions of high recombination, including a recombination hotspot within TAP2 [97], and in a LD block that includes few other HLA genes. These data suggest that the signal of balancing selection at rs2071473 is distinct from other signals of balancing selection in the HLA region, but it is difficult to disentangle these signals given the relatively strong linkage across the HLA region. Thus it is possible that the C/T polymorphism at rs2071473 is not itself under balancing selection and is linked to the balanced site.

If the target of balancing selection is rs2071473, what selective forces are acting to maintain the ancestral and derived alleles? Balancing selection is generally attributed to heterozygote advantage (overdominance), frequency dependent selection, or antagonistic pleiotropy [98], which are all probable evolutionary scenarios to explain maintenance of the ancestral and derived alleles of rs2071473. Intriguingly, GWA studies have found rs2071473 genotype is associated with ulcerative colitis [99], Crohn’s disease [100], and sarcoidosis [101,102] in addition to fecundability. For example, the ancestral T allele is significantly associated with ulcerative colitis [99] and shorter time to pregnancy whereas the derived C allele is significantly associated with Crohn’s disease [100] and longer time to pregnancy suggesting theses alleles may be maintained antagonistic pleiotropy.

## Methods

### rs2071473 is a multi-tissue eQTL for TAP2, HLA-DOB, and HLA-DRB6

We replicated the association between the T/C polymorphism at rs2071473 and ***TAP2*** expression levels using GTEx Analysis Release V6 (dbGaP Accession phs000424.v6.p1) data for 35 tissues including the uterus [22,23]. We also used GTEx data to identify other genes for which rs2071473 was an eQTL. Briefly, we queried the GTEx database using the ‘Single tissue eQTLs search form’ for SNP rs2071473 (http://www.gtexportal.org/home/eqtls/bySnp).

### *TAP2* is expressed by decidual stromal cells at the maternal-fetal interface

To determine the cell-type localization of TAP2 in the endometrium, we used data from the Human Protein Atlas immunohistochemistry collection (www.proteinatlas.org)[24] for endometrium, placenta, and decidua. We examined the expression of ***TAP2*** in the endometrium across the menstrual cycle [25], in mid-secretory phase endometrial biopsies from women not taking hormonal contraceptives (n=11), and women using either the progestin-based contraceptives depot medroxyprogesterone acetate (DMPA) or levonorgestrel intrauterine system (LNG-IUS) for at least 6 months [26], and in DSCs treated with a PGR-specific siRNA or a non-targeted siRNA [27] using previously generated microarray expression data. These microarray datasets were analyzed with the GEO2R analysis package (http://www.ncbi.nlm.nih.gov/geo/geo2r/), which implements the GEOquery [28] and limma R packages [29,30] from the Bioconductor project to quantify differential gene expression. We also examined ***TAP2*** expression in RNA-Seq data previously generated from ESFs treated with control media or differentiated (decidualized) with cAMP/MPA into DSCs [31,32].

### Cell culture and luciferase assays

#### Cell culture

Endometrial stromal fibroblasts (ATCC CRL-4003) immortalized with telomerase were maintained in phenol red free DMEM (Gibco) supplemented with 10% charcoal stripped fetal bovine serum (CSFBS; Gibco), 1× ITS (Gibco), 1% sodium pyruvate (Gibco), and 1% L-glutamine (Gibco). Decidualization was induced using DMEM with phenol red (Gibco), 2% CSFBS (Gibco), 1% sodium pyruvate (Gibco), 0.5mM 8-Br-cAMP (Sigma), and 1μM medroxyprogesterone acetate (Sigma).

#### Plasmid Transfection and Luciferase Assays

A 1000bp region spanning 50bp upstream of rs2071732 to the 3’-end of the PGR ChIP-Seq peak was cloned into the pGL3-Basic luciferase vector (Promega), once with the T allele and once with the C allele (Genscript). A pGL3-Basic plasmid without the 1kb rs2071473 insert was used as a negative expression control. DDIT3 and PGR expression plasmids were also obtained from Genscript. Confluent ESFs in 96 well plates in 80μl of Opti-MEM (Gibco) were transfected with 100ng of the luciferase plasmid, 100ng of DDIT3 and/or PGR as needed, and 10ng of pRL-null with 0.1 μl PLUS reagent (Invitrogen) and 0.25μl of Lipofectamine LTX (Invitrogen) in 20μl Opti-MEM. The cells incubated in the transfection mixture for 6hrs and the media was replaced with the phenol red free maintenance media overnight. Decidualization was then induced by incubating the cells in the decidualization media for 48hrs. After decidualization, Dual Luciferase Reporter Assays (Promega) were started by incubating the cells for 15mins in 20μl of 1x passive lysis buffer. Luciferase and renilla expression were then measured using the Glomax multi+ detection system (Promega). Luciferase expression values were standardized by the renilla expression values and background expression values as determined by pGL3-Basic expression.

### The rs2071473 C allele is derived in humans and segregating at intermediate to high frequencies across multiple human populations

To reconstruct the evolutionary history of the T/C polymorphism we used a region spanning 50bp upstream and downstream of rs2071473 from hg19 (chr6:32814778-32814878) as a query sequence to BLAT search the chimpanzee (CHIMP2.1.4), gorilla (gorGor3.1), organutan (PPYG2), gibbon, (Nleu1.0), rhesus monkey (MMUL_1), hamadryas baboon (Pham_1.0), olive baboon (Panu_2.0), vervet monkey (ChlSab1.0), marmoset (C_jacchus3.2.1), Bolivian squirrel monkey (SalBol1.0), tarsier (tarSyr1), mouse lemur (micMur1), and galago (OtoGar3) genomes. For all other non-human species we used the same 101 bp region as a query for SRA-BLAST against high-throughput sequencing reads deposited in SRA. The top scoring 100 reads were assembled into contigs using the ‘Map to reference’ option in Geneious v6.1.2 and the human sequence as a reference. Sequences for the Altai Neanderthal, Denisovan, Ust-Ishim, and two aboriginal Australians were obtained from the ‘Ancient Genome Browser’ (http://www.eva.mpg.de/neandertal/draft-neandertal-genome.html). The frequency of the C/T allele across the Human Genome Diversity Project (HGDP) populations was obtained from the ‘Geography of Genetic Variants Browser’ (http://popgen.uchicago.edu/ggv/).

We inferred ancestral sequences of the 101 bp region using the ancestral sequence reconstruction (ASR) module of the Datamonkey web-sever (http://www.datamonkey.org) [33] which implement joint, marginal, and sampled reconstruction methods [34], the nucleotide alignment of the 101 bp, the best fitting nucleotide substitution model (HKY85), a general discrete model of site-to-site rate variation with 3 rate classes, and the phylogeny shown in **Fig 4A**. All three ASR methods reconstructed the same sequence for the ancestral human sequence at 1.0 support.

### rs2071473 has signatures of balancing in humans and diversifying selection in primates

To infer if there was evidence for positive selection acting on the derived T allele we used an improved version of the EvoNC method [35] that has been modified into a branch-site model [36,37], and implemented in HyPhy v2.22 [37,38]. This method utilizes the MG94HKY85 nucleotide substitution model, which was also the best-fitting nucleotide substitution model for our target non-coding region and neutral rate proxy, and includes 10 replicate likelihood searches. Although this implementation of the EvoNC method that has been modified into a branch-site model capable of identifying positively selected sites in a priori defined lineages using Naive Empirical Bayes (NEB) and Bayes Empirical Bayes (BEB), previous studies have shown that these methods have high type-I and type-II error rates. Therefore, we inferred evidence for without reference to NEB or BEB identified sites and instead relied on a significant likelihood ratio test between the null and alternate models. We analyzed an alignment of the 101 bp region described above, the phylogeny shown in **Fig 4A**, synonymous sites from the flanking *HLA-DOB* and ***TAP2*** genes which were identified from the same primate species using the method described above. Fst, Tajima’s D, and Fay and Wu’s H data for CEU and YRI populations were obtained from the 1000 Genomes Selection Browser 1.0 [103].

## Acknowledgments

This work was funded by a University of Chicago new lab startup package, a Burroughs Wellcome Preterm Birth Initiative grant, and a March of Dimes Transdisciplinary center grant (V.J.L.). The authors thank C. Ober for comments on an earlier version of this manuscript.

